# The MALDI TOF E2/E3 ligase assay as an universal tool for drug discovery in the ubiquitin pathway

**DOI:** 10.1101/224600

**Authors:** Virginia De Cesare, Clare Johnson, Victoria Barlow, James Hastie, Axel Knebel, Matthias Trost

**Affiliations:** MRC Protein Phosphorylation and Ubiquitylation Unit, University of Dundee, Dow St, Dundee, DD1 5EH, Scotland, UK; MRC Protein Phosphorylation and Ubiquitylation Unit Reagents and Services, University of Dundee, Dow St, Dundee, DD1 5EH, Scotland, UK; Institute for Cell and Molecular Biosciences, Newcastle University, Framlington Place, Newcastle-upon-Tyne, NE2 1HH, UK

**Keywords:** Ubiquitin, E3 ligase, E2 enzyme, MALDI TOF, mass spectrometry, drug discovery, high-throughput, assay, MDM2, HOIP, ITCH

## Abstract

In many diseases, components of the ubiquitin system - such as E2/E3 ligases and deubiquitylases - are dysregulated. The ubiquitin system has therefore become an emergent target for the treatment of a number of diseases, including cancer, neurodegeneration and autoimmunity. Despite of the efforts in this field, primary screenings of compound libraries to individuate new potential therapeutic molecules targeting the ubiquitin pathway have been strongly limited by the lack of robust and fast high-throughput assays. Here we report the first label-free high-throughput screening (HTS) assay for ubiquitin E2 conjugating enzymes and E3 ligases based on Matrix-Assisted Laser Desorption/Ionization Time-Of-Flight (MALDI TOF) mass spectrometry. The MALDI TOF E2/E3 assay allows us to test E2 conjugating enzymes and E3 ligases for their ubiquitin transfer activity, to identify E2/E3 active pairs, inhibitor potency and specificity and to screen compound libraries *in vitro* without synthesis of chemical or fluorescent probes. We demonstrate that the MALDI TOF E2/E3 assay is a universal tool for drug discovery screening in the ubiquitin pathway as it is suitable for working with all E3 ligase families and requires a reduced amount of reagents, compared to standard biochemical assays.

## Introduction

Ubiquitylation is a post-translational modification which impacts almost every biological process in the cell. Dysregulation of the ubiquitylation pathway is associated with several diseases, including cancer, neurodegenerative disorders and immunological dysfunctions. Single ubiquitin moieties or polyubiquitin chains are added to the substrate by the combined action of three different classes of enzymes: the E1 activating enzymes, the E2s conjugating enzymes and the E3 ligase enzymes^1^. In the first step, a single ubiquitin molecule is coupled to the active site of an E1 ubiquitin activating enzyme in an ATP-dependent reaction. In the second step, the ubiquitin molecule is transferred from E1 to an E2 ubiquitin conjugating enzyme. In the final step, ubiquitin is transferred to the protein substrate in a process mediated by an E3 ubiquitin ligase, which provides a binding platform for ubiquitin-charged E2 and the substrate. Ubiquitin chain formation is highly specific and regulated by a plethora of different E2 conjugating enzymes and E3 ligases. The human genome encodes two ubiquitin-activating E1, >30 ubiquitin specific E2 and 600–700 E3 ligases^2^. Thus, including about 100 deubiquitylating enzymes, approximately 800 ubiquitin enzymes regulate the dynamic ubiquitylation of a wide range of protein substrates^2^. Within this complexity, E3 ligases are the most diverse class of enzymes in the ubiquitylation pathway as they play a central role in determining the selectivity of ubiquitin-mediated protein degradation and signalling.

E3 ligases have been associated with a number of pathogenic mechanisms. Mutations in the E3 ligases MDM2, BRCA1, TRIMs and Parkin have been linked to multiple cancers and neurodegenerative diseases^3, 4, 5^ and MDM2-p53 interaction inhibitors have already been developed as potential anti-cancer treatment^6^. This highlights the potential of E2 enzymes and E3 ligases as drug targets. Although all E3 ligases are involved in the final step of covalent ubiquitylation of target proteins, they differ in both structure and mechanism and can be classified in three main families depending on the type of E3 ligases promoted ubiquitin-protein ligation and on the presence of characteristic domains. The Really Interesting New Genes (RING) ligases bring the ubiquitin-E2 complex into the molecular vicinity of the substrate and facilitate ubiquitin transfer directly from the E2 enzyme to the substrate protein. In contrast, Homologous to the E6-AP C Terminus family (HECTs) covalently bind the ubiquitin via a cysteine residue in their catalytic HECT domain before shuttling it onto the target molecule. RING between RINGs (RBRs) E3-ligases were shown to use both RING and HECT-like mechanisms were ubiquitin is initially recruited on a RING domain (RING1) then transferred to the substrate through a conserved cysteine residue in a second RING domain. The vast majority of human E3 enzymes belong to the RING family, while only 28 belong to the HECT and 14 to the RBR family of E3 ligases^7^.

Due to the high attractiveness of E2 and E3 ligases as drug targets, a number of drug discovery assays have been published, based on detection by fluorescence^8, 9, 10^, antibodies^11, 12, 13, 14^, tandem ubiquitin-binding entities (TUBEs)^15, 16^, surface plasmon resonance (SPR)^17^ or cellular^18^ and bacterial^19^ two hybrid. However, many of these tools are either too expensive for very high-throughput drug discovery or potentially result in false-positive and false negative hits due to the use of non-physiological E2/E3 ligase substrates. We have addressed this gap by developing the first *in vitro* label-free MALDI TOF mass spectrometry-based approach to screen the activity of E2 and E3 ligases that uses unmodified monoubiquitin as substrate. As a proof of concept, we screened a collection of 1443 FDA approved drugs for inhibitors of a subset of three E3 ligases that are clinically relevant and belong to three different E3 ligase families. The screen shows high reproducibility and robustness and we were able to identify a subset of 15 molecules active against the E3 ligases tested. We validated the most powerful positive hits by determining the IC50 values against their targets, confirming that Bendamustine and Candesartan Cilexitel inhibit HOIP and MDM2, respectively, in *in vitro* conditions.

## Results

### MALDI TOF E2-E3 assay rational and development

E2 and E3 ligase activity results in formation of free or attached polyubiquitin chains, mono-ubiquitylation and/or multiple mono-ubiquitylation of a specific substrate. However, in absence of a specific substrate, most E3 ligases will either produce free polyubiquitin chains or undergo auto-ubiquitylation which is a mechanism thought to be responsible for the regulation of the E3 enzyme itself^20^. Furthermore, there is some evidence that auto-ubiquitylation of E3 ligases is facilitating the recruitment of the E2 ubiquitin-conjugating enzyme^21^. Auto-ubiquitylation assays or free polyubiquitin chain production have been widely used to assess the E3 ligase potential of a protein^20, 22^. We used this property of E2 and E3 ligases to design a MALDI TOF mass spectrometry-based high-throughput screening (HTS) method that allowed the reliable determination of activities of E2 and E3 ligase pairs by measuring the depleting intensity of mono-ubiquitin in the assay as a readout.

As proof-of-concept we used three E3 ligases belonging to different E3 families and representative of all the currently known ubiquitylation mechanisms. MDM2 is an RING-type E3 ligase which controls the stability of the transcription factor p53, a key tumour suppressor that is often found mutated in human cancers^23, 24^. ITCH belongs to the HECT domain-containing E3 ligase family involved in the regulation of immunological response and cancer development^25, 26, 27^. Finally, HOIP, a RBR E3 ubiquitin ligase and member of the LUBAC (linear ubiquitin chain assembly complex). As part of the LUBAC complex, HOIP is involved in the regulation of important cellular signalling pathways that control innate immunity and inflammation through nuclear factor (NF)-kB activation and protection against tumour necrosis factor α (TNFα)-induced apoptosis^28, 29^. HOIP is the only known E3 ligase generating linear ubiquitin chains^30^. Because of that, fluorescent assays using C-terminally or N-terminally labelled ubiquitin species cannot be used to form linear chains.

To determine MDM2, ITCH and HOIP auto-ubiquitylation reaction rate and the linearity range we followed the consumption of mono-ubiquitin over time with increasing starting amount of mono-ubiquitin. We matched MDM2, ITCH and HOIP with E2 conjugating enzymes as reported in literature: MDM2 and ITCH were incubated with E2D1 (UbcH5a)^31^ while HOIP was used in combination with UBE2L3 (UbcH7)^28^. Briefly, the *in vitro* ubiquitylation reaction consisted of 1 mM ATP, 12.5, 6.25 and 3.125 μM ubiquitin (Ub), 50 nM E1, 250 nM E2 and 250 or 500 nM E3 ligase enzyme at 37°C for 30 min in a total volume of 5 μl (**Figure 1A**). Reactions were started by addition of Ub and terminated by addition of 2.5 μl of 10% (v/v) trifluoroacetic acid (TFA). 1.05 μl of each reaction was then spiked with 300 nl (4 μM) of ^15^N-labelled ubiquitin and 1.2 μl of 2,5-dihydroxyacetophenone (DHAP) matrix and 250 nL of this solution was spotted onto a 1,536 microtiter plate MALDI anchor target using a nanoliter dispensing robot. The samples were analysed by high mass accuracy MALDI TOF MS in reflector positive ion mode on a rapifleX MALDI TOF mass spectrometer.

**Figure 1:**
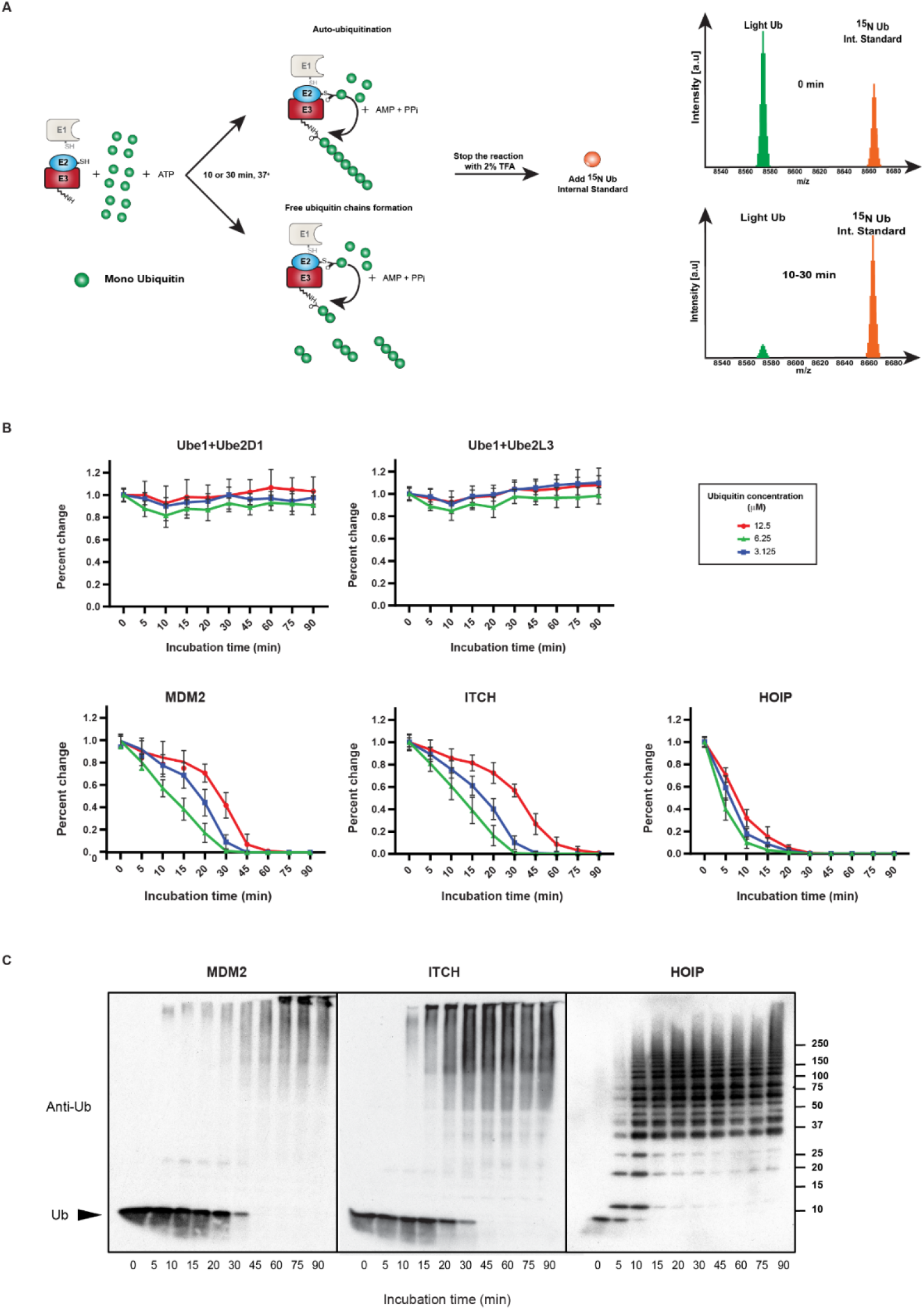
The MALDI TOF E2/E3 ligase assay. (a) Workflow of the MALDI TOF E2/E3 assay. Each of the three E3 ligases were incubated with their E2 partner with different concentrations of monoubiquitin (12.5, 6.25 and 3.125 μM) at 37°C. Reactions were stopped by addition of 2.5 μL 10% TFA at different time points. 1.05 μl of reaction aliquots were mixed with 150 nl of 1.5 μM ^15^N Ubiquitin as internal standard. Subsequently, the analytes were mixed with 2,5 DHAP matrix and spotted onto a 1,536 AnchorChip MALDI target (Bruker Daltonics). Data analysis was performed using FlexAnalysis. (b) E2/E3 ligase reactions are linear. Linearity is determined by monoubiquitin consumption over time. Only at very high and low concentrations of ubiquitin, the reaction is not linear. Data points have been normalized to determine reaction linearity. (c) Western blots of *in vitro* reactions of the three E3 ligases showing increased ubiquitin chain formation over time.

Importantly, the use of ^15^N ubiquitin as internal standard allowed us not only to avoid spot-to-spot and shot-to-shot variability in MALDI ionization^32^, but also it allowed us to keep track of the amount of mono-ubiquitin “consumed” during the assay. Overall, this setup allowed us to achieve very high precision, accuracy and reproducibility of measurements.

In our experimental conditions, Ub-consumption relied on the presence on an E3 ligase as we did not observe a significant reduction in the ubiquitin level within the negative controls (**Figure 1B**), where only E1 activating enzyme and E2 conjugating enzyme were present. We found that the optimal concentrations for the enzymes used were in the nanomolar range (250 nM for HOIP, 500 nM for ITCH and 500 nM for MDM2). As expected, we observed that Ub consumption is dose and enzyme dependent (**Figure 1B, Supplementary Fig. 1**). Reaction rates were related to Ub concentration (**Supplementary Table 1**) and different enzymes showed different rates of ubiquitin consumption (**Figure 1B**).

The well-established E2-E3 auto-ubiquitylation assays followed by SDS-Page and western blot analysis provided similar results and we observed that the time dependent disappearance of Ub is comparable using both techniques (**Figure 1C**). Moreover, while substrate and enzyme concentrations are comparable to Western Blot based approaches, the reaction volume (5 μl) is smaller than most of the antibody-based approaches currently reported in literature^33^.

### Determining *in vitro* activities of E2 enzymes

E2 ubiquitin-conjugating enzymes are the central players in the ubiquitin cascade^34^. The human genome encodes ~40 E2 conjugating enzymes, of which about 30 conjugate ubiquitin directly while others conjugate small Ubiquitin-like proteins such as SUMO1 and NEDD8^35^. E2 enzymes are involved in every step of the ubiquitin chain formation pathway, from transferring the ubiquitin to mediating the switch from ubiquitin chain initiation to elongation and defining the type of chain linkage. Connecting ubiquitin molecules in a defined manner by modifying specific Lys residues with ubiquitin is another intrinsic property of many E2 enzymes. Early studies showed that at high concentrations, E2 enzymes can synthesize ubiquitin chains of a distinct linkage or undergo auto-ubiquitylation even in the absence of an E3^36^, albeit at lower transfer rates^34^. This characteristic has been exploited for the generation of large amounts of different ubiquitin chain types *in vitro*^37^.

As control for E2 enzyme mono or multi-ubiquitylation or E2-dependent ubiquitin chain assembly, we firstly assessed which E2 conjugating enzymes in our panel were able to consume ubiquitin even in absence of a partner E3 ligase. Utilizing the MALDI TOF E2-E3 assay, we systematically tested 27 recombinantly expressed E2 conjugating enzymes (**Supplementary Table 2**) for their ability to process ubiquitin either by the formation of polyubiquitin chains or by auto-ubiquitylation at different concentrations (250 nM, 500 nM and 1 μM). We found that the UBE2Q1 and UBE2Q2 were able to consume ubiquitin even in absence of a specific E3 ligase at 250 nM after 45 min incubation time and almost completely exhausting the starting ubiquitin amount after 2h of incubation (**Figure 2**). UBE2O and UBE2S are able to consume ubiquitin when present at a starting concentration of 500 nM with consumption being evident from 90 min onwards. Interestingly, UBE2Q1, UBE2Q2 and UBE2O^38, 39, 40^ are E2 conjugating enzymes characterized by an unusually high molecular mass compared to other E2 enzymes: in particular UBE2O has been reported as an E2-E3 hybrid which might explain its ability to form ubiquitin chains in absence of an E3 ligase. Most of the E2 conjugating enzymes showed ubiquitin-consuming activity once their concentration was increased to 1 μM. Interestingly, UBE2D1 and UBE2L3 do not show any ligase activity even at 1 μM, making these E2 conjugating enzymes the perfect candidates for inhibitor screening of E3 ligases as all ubiquitin consuming activity in an assay will be down to E3 activity. Our results demonstrate that the E2-E3 MALDI TOF assay has the potential to be employed for measuring E2 *in vitro* activity and therefore can be employed for screening inhibitors against those E2 enzymes that possess intrinsic ubiquitin ligase activity *in vitro*.

**Figure 2:**
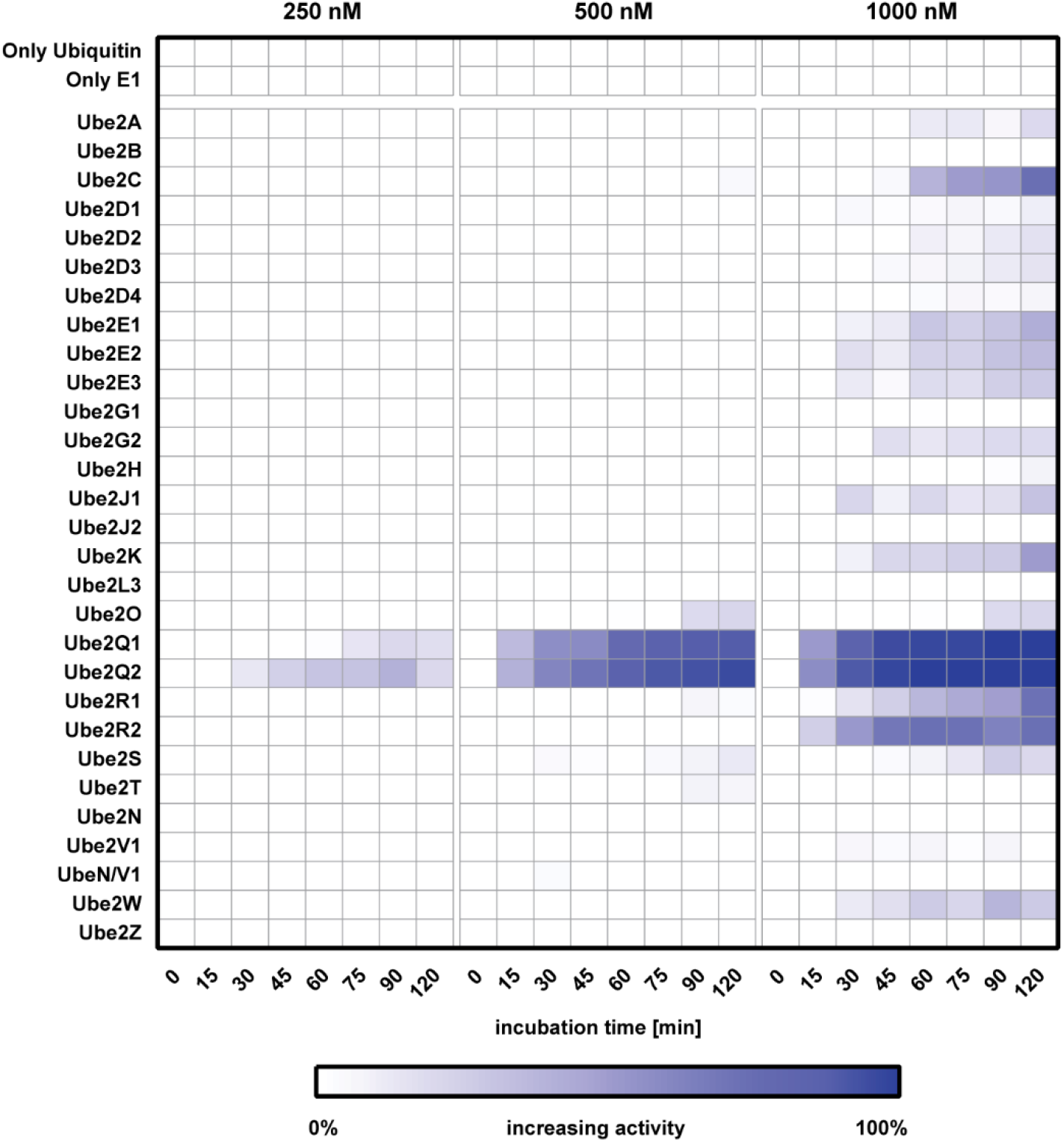
Characterizing E2 enzyme *in vitro* activity. 27 E2 enzymes were incubated at three different concentrations (250-500-1000 nM) with the E1 UBE1 and 6.125 μM ubiquitin at 37°C. Reactions were stopped at the indicated time points with 2% TFA final and analysed by MALDI TOF mass spectrometry. Ligase activity was calculated considering T0 as 0% and the complete disappearance of mono-ubiquitin from the windows signal as 100% activity.

### Determining E2/E3 active pairs

Any given E3 ligase cooperates with specific E2 enzymes *in vivo*. However, it is still difficult to predict which E2/E3 enzyme pair would be functional. Determining E2/E3 specificity is paramount to set up *in vitro* ubiquitylation assays and to perform inhibitor screens against E3 ligases. Using the E2/E3 MALDI TOF assay, we investigated the activity of MDM2, ITCH and HOIP throughout eight time points when incubated with any of 27 ubiquitin E2 enzymes, covering the majority of the reported classes/families (**Figure 3**). We arbitrarily defined “fully active” pairs any E2-E3 couple that completely depleted the mono-ubiquitin starting amount after 2 hours incubation time. The UBE2D family is reported in literature as being able to productively interact with MDM2^41^, ITCH^33^ and HOIP^42^. Our data showed that UBE2D1 and UBE2D2, also known as UBCH5a and UBCH5b, were fully active with all the E3 ligases under investigation confirming the promiscuous activity of this class of E2 enzymes that was previously reported in the literature^33, 41, 42^. The UBE2E family was only partially active against the E3s ligases of interest. The well-characterized human E2L3 (or UBCH7) showed activity with HOIP and ITCH, confirming the already reported UBCH7 ability of functioning with both HECT and RBR E3 families^43, 44^. Taken together these results demonstrate that the E2/E3 MALDI TOF assay is suitable for determining E2 specificity towards their cognate E3 enzymes in a high-throughput fashion.

**Figure 3:**
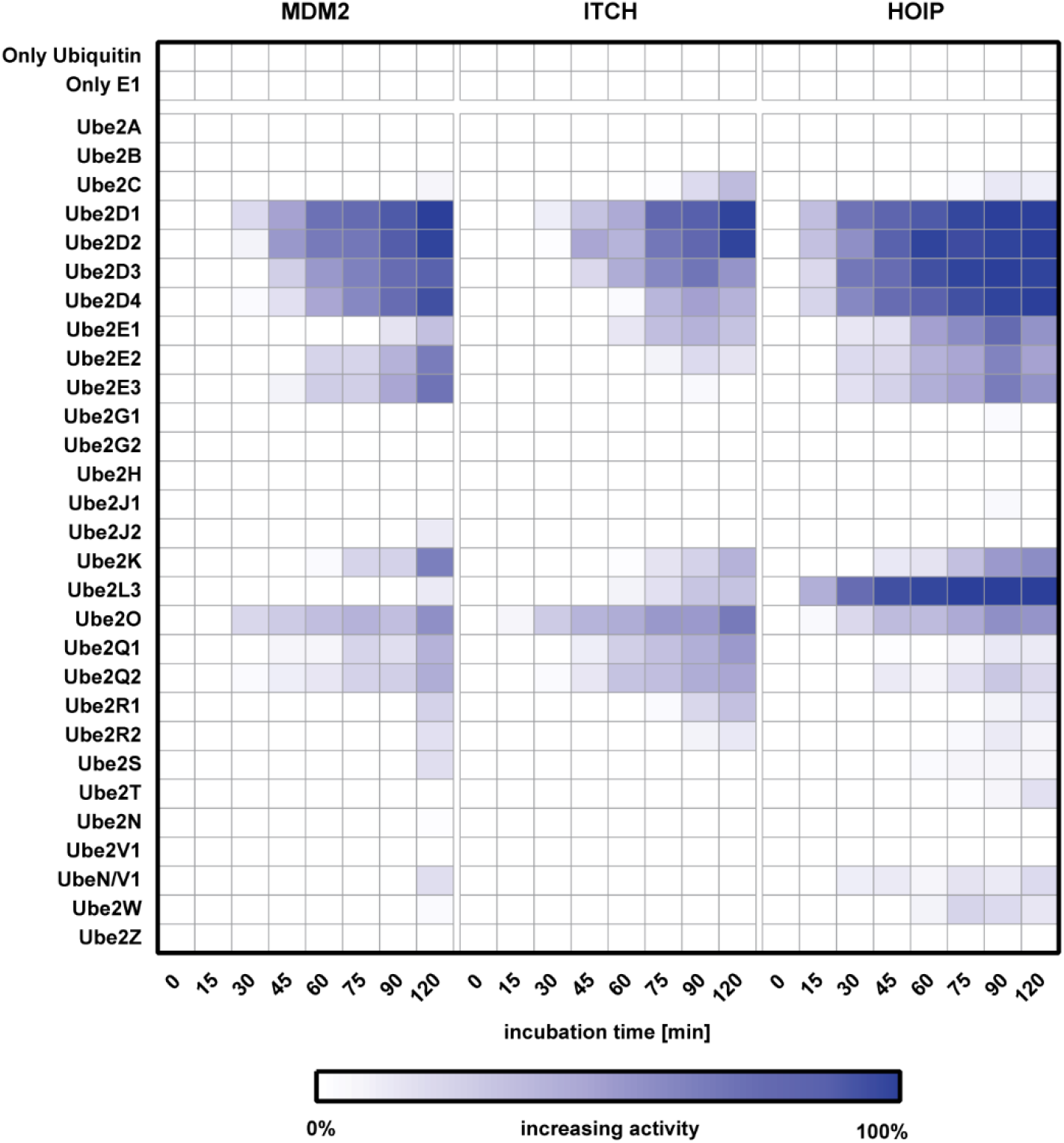
Characterizing E2/E3 pair activities. E1-E2 enzymes and E3 ligases were incubated in duplicate with 6.125 μM ubiquitin at 37°C for the time indicated. Reactions were stopped at the indicated time points with 2% TFA final and analysed by MALDI TOF mass spectrometry. Ligase activity was calculated considering T0 as 0% and the complete disappearance of mono-ubiquitin signal from the mass window as 100% activity.

### Assessing potency and selectivity of E2/E3 inhibitors

We next evaluated whether the MALDI TOF E2/E3 assay had potential to assess the potency and selectivity of E2/E3 inhibitors. As proof-of-concept, we tested 5 inhibitors that had previously been reported to inhibit E1, E2 or E3 ligases: PYR41^45^, Bay117082^46^, Gliotoxin^47^, Nutlin3A^48^ and Clomipramine^49^. We also tested PR619^50^ a broad spectrum, alkylating DUB inhibitor that we hypothesised would also inhibit other enzymes with active site cysteines, such as E1/E2 enzymes and E3 ligases. PYR41 is a specific and cell permeable inhibitor of E1 ubiquitin loading but does not directly affect E2 activity^45^. Bay117082, initially described as inhibitor of nuclear factor κB (NF-κB) phosphorylation, has been shown to inactivate the E2-conjugating enzymes Ubc13 (UBE2N) and UBCH7 (UBE2L3) as well as the E3 ligase LUBAC (of which HOIP is part)^46^. Gliotoxin is a fungal metabolite identified as a selective inhibitor of HOIP through a FRET-based HTS assay^47^. Nutlin 3A is a MDM2-p53 interaction inhibitor, able to displace p53 from MDM2 with an IC50 in the 100 to 300 nM range^51^. However Nutlins are not reported to be able to inhibit MDM2 auto-ubiquitylation. Clomipramine is a compound reported as able to block ITCH Ub transthiolation in an irreversible manner, achieving complete inhibition at 0.8 mM^49^. Performing IC50 inhibition curves using the MALDI TOF E2/E3 assay (**Figure 4, Supplementary Table 3**), we could show that PYR41 inhibited both HOIP and ITCH at IC50s of 5.4 μM and 11.3 μM, respectively. Bay117082 also strongly inhibited HOIP and ITCH with an IC50 of 2.9 μM and 25.9 μM, respectively. It is important to underline that the IC50 calculation maybe be affected by the content in DTT/TCEP and or β-mercaptoethanol as these compound might also react with compounds that are highly reactive towards thiol groups. Gliotoxin showed an IC50 of 2.8 μM against HOIP and 30.61 μM against ITCH but it did not inhibit MDM2. As expected, Nutlin 3A did not show any inhibitory activity towards MDM2, ITCH or HOIP as it was designed as an interaction inhibitor. We found that Clomipramine acted as a weak inhibitor for ITCH with an IC50 of 503.4 μM but it did not affect MDM2 or HOIP. Our results demonstrate that the MALDI TOF E2-E3 assay is suitable for comparing drug potency through IC50 determination.

**Figure 4:**
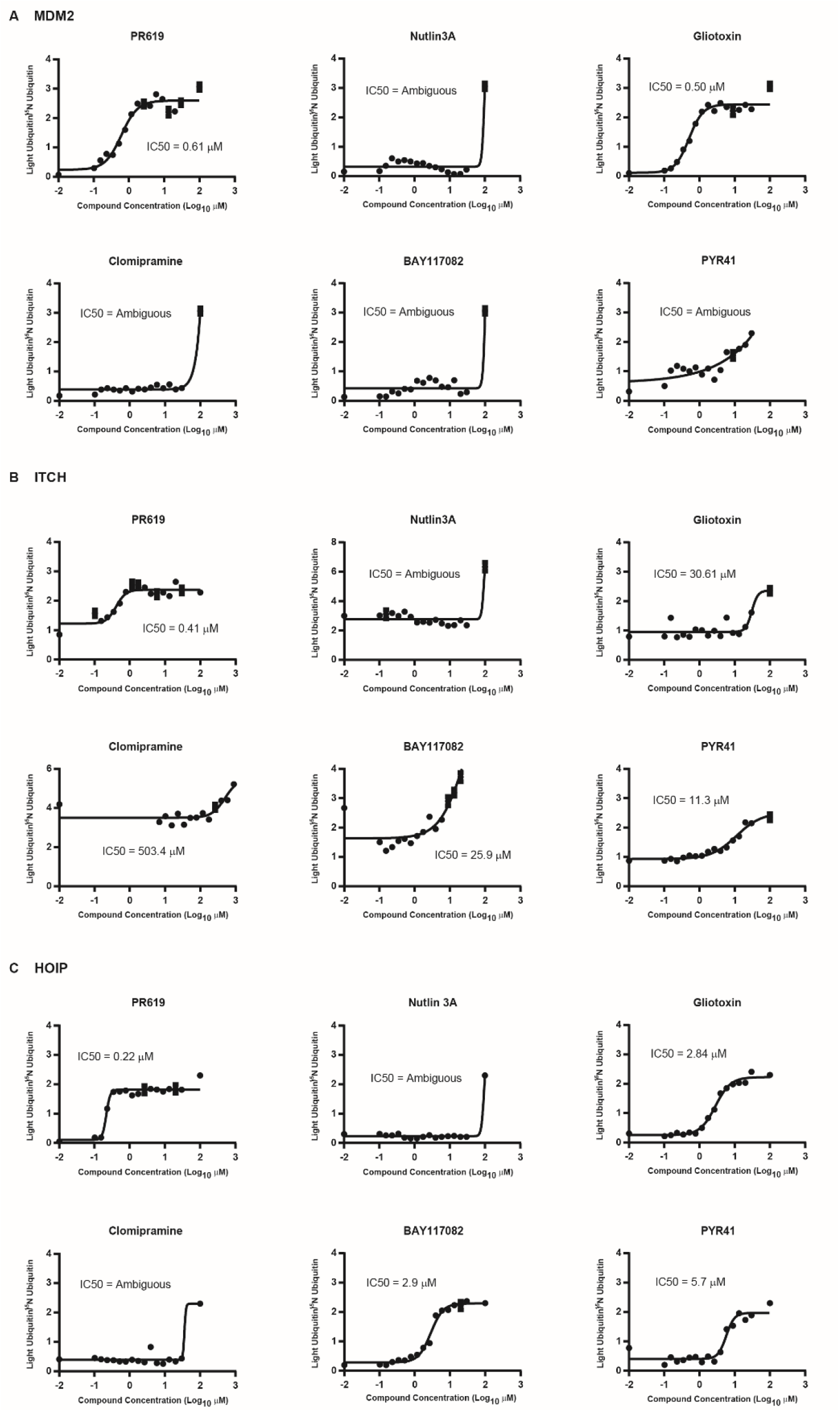
IC50 analyses of six Inhibitors for selected E3 ligases. IC50 determination for six described E3 ligase inhibitors for MDM2, ITCH and HOIP. Small inhibitor compounds were pre-incubated for 30 min at different concentrations (0–100 μM). Ubiquitin was added and incubated for a range of time depending on the E3 (usually 30 or 40 minutes). For statistical analysis, Prism GraphPad software was used with a built-in analysis, nonlinear regression (curve-fit), variable slope (four parameters) curve to determine IC50 values.

### E2/E3 assay by MALDI TOF is suitable for HTS

Having established that the E2/E3 MALDI TOF assay can be used to assess the specificity and potency of inhibitors, we explored its suitability for high-throughput screening. It is important to underline that, because of the nature of the assay, inhibitors of either E1 or E2, which both contain active site cysteines ^47, 49^ that form thioester intermediates with ubiquitin may be identified as hits. We tested a library of 1430 FDA approved compounds from various commercial suppliers with validated biological and pharmacological activities at 10 μM final. None of the compounds present in the library are known for specifically targeting MDM2, ITCH or HOIP. The assay was performed supplying ATP in excess (1 mM) to reduce the likelihood of identifying ATP analogues as inhibitors of these enzymes.

The screens against the three different E3 ligases, expressed as percentage effect (**Figure 5, Supplementary Figure 2**) exhibited robust Z’ (Z-prime) scores > 0.5 (**Supplementary Table 4**). These scores provide a measure for the suitability of screening assays; HTS assays that provide Z’ scores > 0.5 are generally considered robust. We defined as a positive hit a compound whose potency ranked above the 50% residual activity threshold. Overall, we identified 9 compounds reporting inhibition rates >50% against the E3 ligases of interest. Candesartan Cilexetil was the only compound able to inhibit MDM2 activity more than 50% (see **Table 1**). With regard to HOIP screening, six compounds were identified as potential inhibitors: Bendamustine, Moclobemide, Ebselen, Cefatrizine, Fluconazole and Pyrazinamide. The ITCH inhibitor screening identified two positive hits: Hexachlorophene and Ethacrynic Acid. Hexachlorophene is an organochloride compound once widely used as a disinfectant. It acts as an alkylating agent thus resulting in the wide and not specific inhibition of E3 ligases: this explains why this compound results as a weak inhibitor of MDM2 as well. Ethacrynic acid is a diuretic compound^52^ and a potent inhibitor of glutathione S-transferase with intrinsic chemical reactivity toward sulfhydryl groups^53^ that might explains its ranking as positive hit in our assay when tested against both MDM2 and ITCH.

**Table 1.**
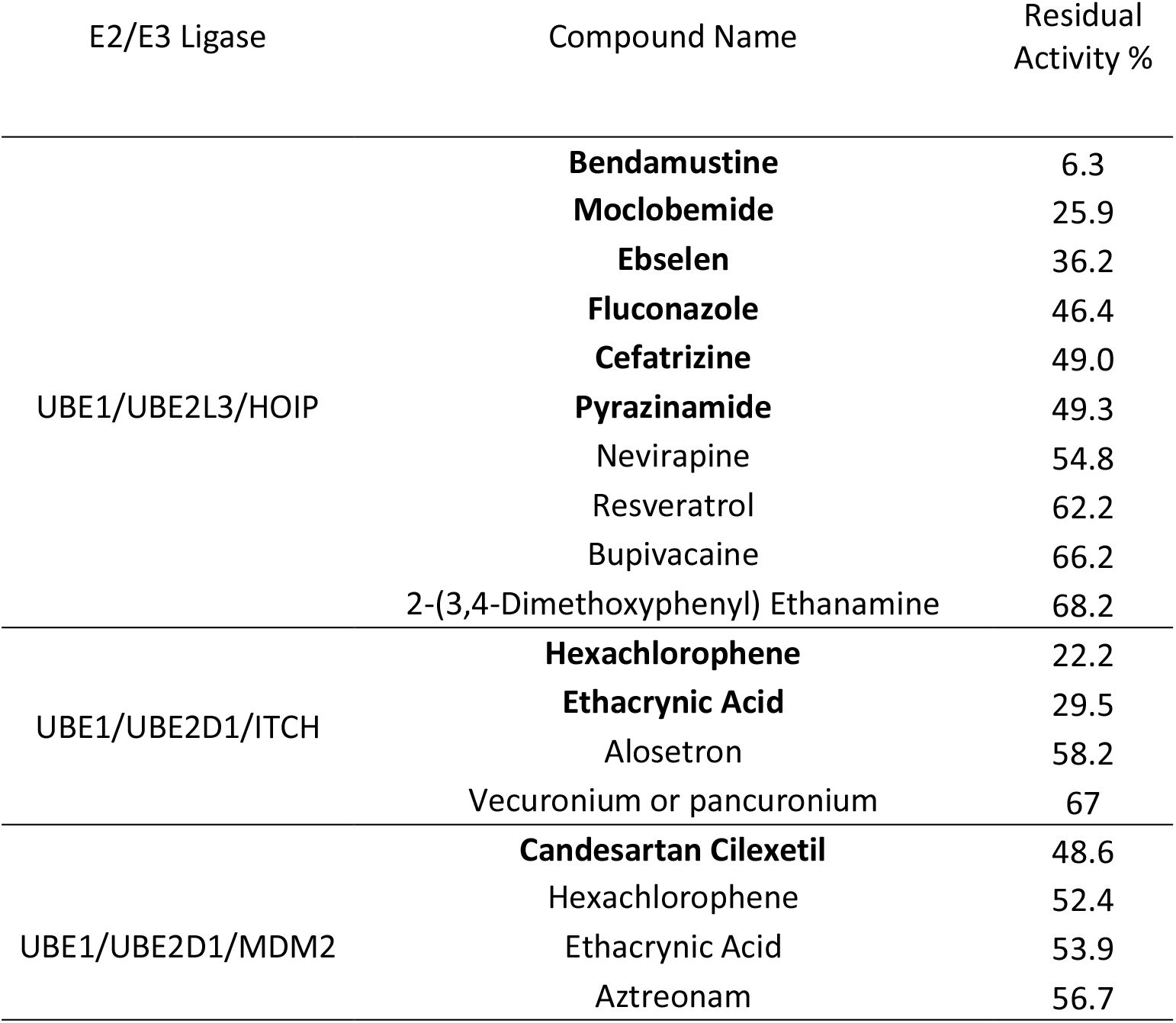
Positive hits identified by E2/E3 MALDI TOF assay

Overall, our results demonstrate that the E2/E3 MALDI TOF assay can be employed to screen large compound libraries against E3 ligases belonging to different families for the identification of new inhibitors.

**Figure 5:**
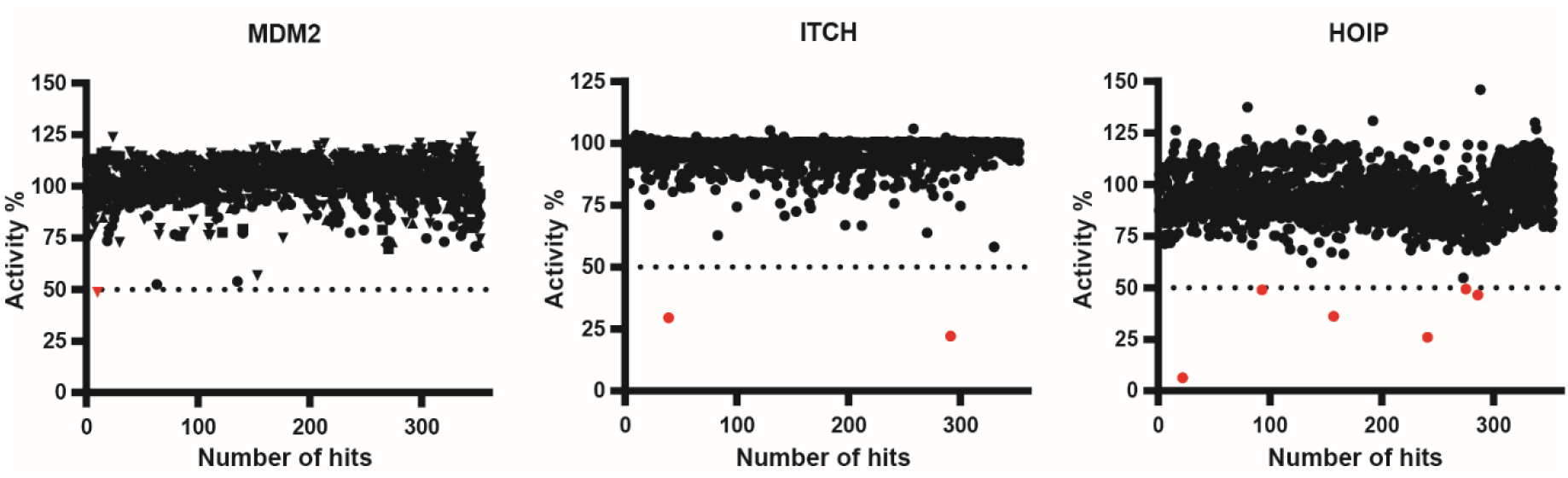
High-throughput E2/E3 MALDI TOF screen. 1430 compounds from various commercial suppliers were tested at a final concentration of 10 μM each against MDM2, ITCH and HOIP. The uninhibited control contained 5 nl DMSO but no compound, whereas the inhibited control had been inactivated by pre-treatment with 2.0 % TFA. Compounds inhibiting reactions >50% are marked in red.

### Validation of positive hits

T o validate the results obtained from the HTS we performed IC50 determination of compounds with the highest inhibitor potency in the single point screening. Candesartan Cilexitel, an angiotensin II receptor antagonist, inhibited MDM2 with an IC50 of 8.9 μM (**Figure 6A**). Best hit was Bendamustine, a nitrogen mustard used in the treatment of chronic lymphocytic leukemia and lymphomas. It belongs to the family of alkylating agents. Bendamustine ranked as the compound with the highest inhibition score against HOIP while it did not significantly affect MDM2 and ITCH activities. We confirmed that Bendamustine selectively inhibited HOIP at an IC50 = 6.4 μM while ITCH and MDM2 showed considerably higher IC50s (113 and 76.8 μM respectively) (**Figure 6B**). Bendamustine retained its inhibition power when HOIP was paired with UBE2D1 as conjugating enzyme (**Supplementary Figure 3**) suggesting that the compound binds preferentially to HOIP. This shows that the MALDI TOF E2/E3 ligase assay can be used to identify selective inhibitors from a high-throughput screen.

**Figure 6:**
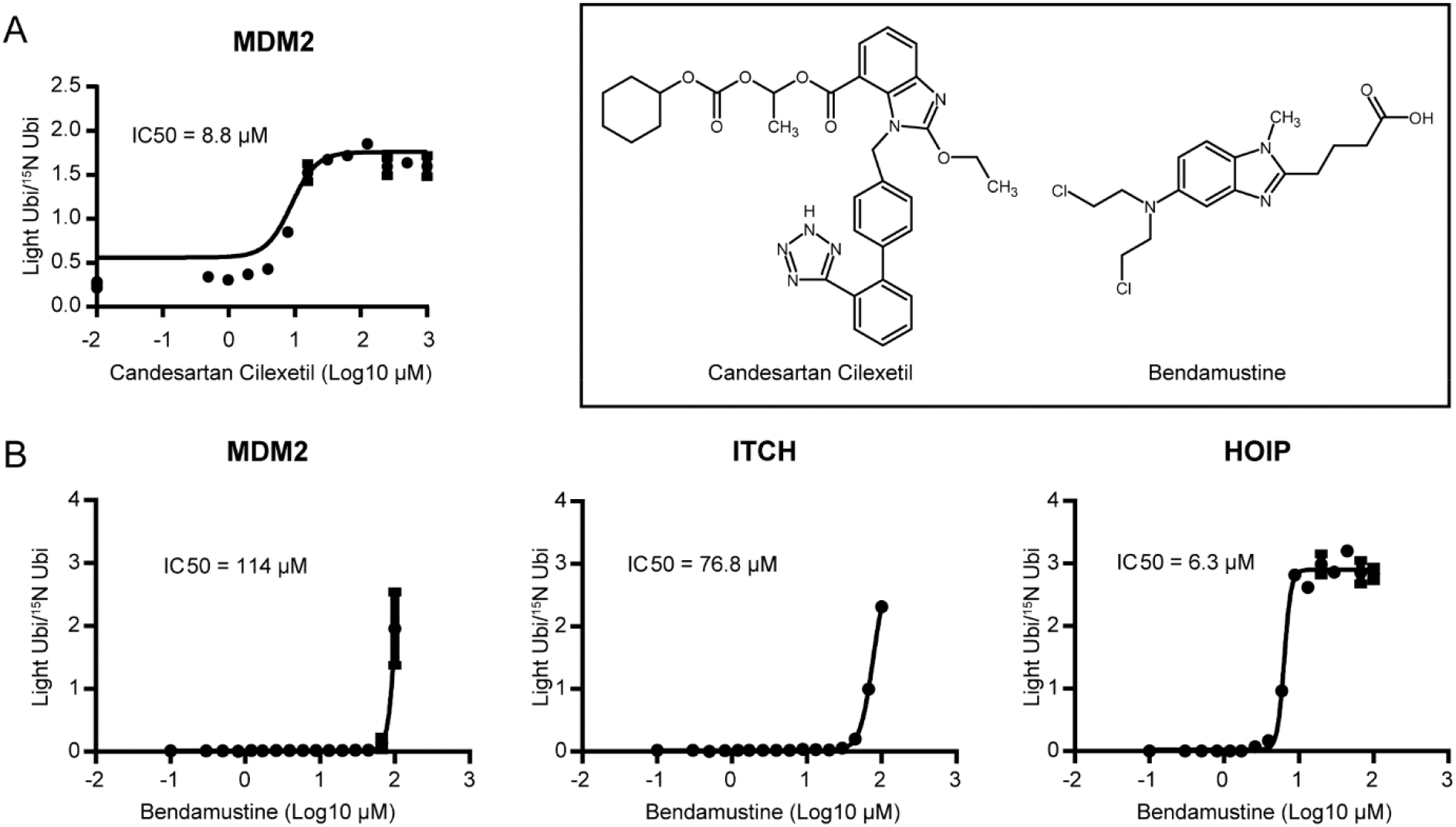
Validation of hits. (A) Candesartan Cilexetil shows an IC50 of 8.8 μM against MDM2. (B) Bendamustine shows some specificity for HOIP (IC50=6.3 μM) while inhibiting MDM2 (IC50=114 μM) and ITCH (IC50=76.8 μM) at higher concentrations.

## Discussion

The ubiquitin system has in recent years become an exciting area for drug discovery^54^ as multiple enzymatic steps within the ubiquitylation process are druggable. The potential of targeting the ubiquitin-proteasome pathway was first demonstrated in 2003 by the approval of the proteasome inhibitor Bortezomib (Velcade; Millennium Pharmaceuticals) for use in multiple myeloma. While proteasome inhibition is a broad intervention affecting general survivability, E3 ubiquitin ligases and deubiquitylases (DUBs)^32^ represent the most specific points of intervention for therapeutic tools as they specifically regulate ubiquitylation rate of specific substrates. For example, Nutlins, cis-imidazoline analogues able to inhibit the interaction between MDM2 and tumour suppressor p53, have recently entered early clinical trials for the treatment of blood cancers^55^. The small number of drugs targeting E3 ligases currently on the market is partly due to the lack of suitable high-throughput assays for drug discovery screening. Traditionally, screening for inhibitors of ubiquitin ligases and deubiquitylases has been performed using different fluorescence-based formats in high-throughput and ELISA, SDS-PAGE and Western blotting in low-throughput. These approaches show a number of limitations. ELISA and SDS-PAGE based approaches are time consuming and low-throughput by nature and therefore mostly incompatible with HTS. The applicability of fluorescence-based techniques such as FRET is dependent on being able to get FRET donors and acceptors in the right distance and the fluorescent label might affect inhibitor binding. To address these issues, we have developed a sensitive and fast assay to quantify *in vitro* E2/E3 enzyme activity using MALDI TOF MS. It builds on our DUB MALDI TOF assay^32^, which has enabled us to screen successfully for a number of selective DUB inhibitors^56, 57, 58^, and adds to the increasing number of drug discovery assays utilising label-free high-throughput MALDI TOF MS. Apart from E2/E3 enzymes and DUBs^32^, high-throughput MALDI TOF MS has now successfully been used for drug discovery screening of protein kinases^59^, histone demethylases and acetylcholinesterases^60^ as well as histone lysine methyltransferases^61^.

Unlike other current assays, all these label-free MALDI TOF MS methods use unmodified substrates, such as mono-ubiquitin. The advantages compared to fluorescence or antibody-based high-throughput assays is the ability to work with enzymes without the previous development of specific chemical/fluorescent probes as well as the reduced consumable costs for the assay as no antibodies are required. Moreover, because of the sensitivity of current MALDI TOF mass spectrometers, all enzymes are usually kept at low concentrations thereby significantly reducing the amounts and cost per assay.

In the context of E3 ligase drug discovery, it is critical to identify the appropriate E2/E3 substrate pairing to ensure the development and use of the most physiologically relevant screening assay. There have been many reports of limited E2/E3 activity profiling with a small number of E2 and E3 enzymes using ELISA-based assays, structural-based yeast two-hybrid assays and Western blot^33, 41, 42^. All of these approaches are time consuming, require large amounts of reagents and are difficult to adapt for HTS. We have successfully used our E2/E3 MALDI TOF assay to identify active E2/E3 pairings which could then be further characterized using our HTS screen. The “E2 scan” was quickly and easily adapted, collecting data of three E3 enzymes against 29 E2 enzymes at eight time-points in one single experiment. Moreover, after identification of the right E2/E3 pairs, we applied the MALDI TOF E2/E3 assay to determine inhibition rates and the IC50 of small molecule inhibitors. In a proof-of-concept study, we performed a HTS for inhibitors of three E3 ligases. The MALDI TOF analysis speed of 1.3 seconds per sample (~35 minutes per 1536 well plate) and low sample volumes (reaction volume 5 μL / MALDI deposition 250 nL) make the E2/E3 MALDI TOF assay comparable to other fluorescence/chemical probe based technologies. Automatic sample preparation, MALDI TOF plate spotting and data collection allowed us to quickly analyse thousands of compounds through the use of 1536-sample targets. The assay successfully identified Bendamustine as a nnovel small molecule inhibitor for HOIP, an attractive drug target for both inflammatory disease and cancer^30, 62, 63, 64^. Bendamustine, a nitrogen mustard, shows likely very high reactivity against a range of targets in the cell including its intended target DNA. However, it is surprising that it shows a 12-fold and 18-fold higher activity against HOIP than against ITCH and MDM2, respectively, suggesting that there is possibly a structural effect and some selectivity can be reached between different E3 ligases. As changes of the E2 enzyme did not change the inhibition of HOIP by Bendamustine, it also shows that E1 and E2 conjugating enzymes were not affected. While this is just a proof-of-concept study characterising E2/E3 activity and identifying inhibitors in an *in vitro* system, follow-up studies will need to verify results in cellular and ultimately *in vivo* models.

In conclusion, we present here a novel screening method to assay E2/E3 activity with high sensitivity, reproducibility and reliability, which is able to carry out precise quantified measurements at a rate of ~1 s per sample spot. Using physiological substrates, we showed proof-of concept for three E3 ligases that are attractive drug targets. Considering the speed, low consumable costs and the simplicity of the assay, the MALDI TOF E3 ligase assay will serve as a sensitive and fast tool for screening for E3 ligase inhibitors.

## Materials and Methods

### Materials

Ubiquitin monomer, BSA, Tris, DTT, Clomipramine and Gliotoxin were purchased from Sigma-Aldrich. MALDI TOF MS materials (targets, matrix and protein calibration mixture) were purchased from Bruker Daltonics (Bremen, Germany). PYR41 and Nutlin3A compounds were kindly provided by Sara Buhrlage, PhD (Dana-Farber Cancer Institute).

### E1, E2, ITCH and HOIP E3 enzyme expression and purification

^15^N-labelled ubiquitin was produced as described in Ritorto et al^32^. Human recombinant 6His-tagged UBE1 was expressed in and purified from Sf21 cells using standard protocols. Human E2s were all expressed as 6His-tagged fusion proteins in BL21 cells and purified via their tags using standard protocols. Briefly, BL21 DE3 codon plus cells were transformed with the appropriate constructs (see table below), colonies were picked for overnight cultures, which were used to inoculate 6 x 1L LB medium supplemented with antibiotics. The cells were grown in Infors incubators, whirling at 200 rpm until the OD_600_ reached 0.5 – 0.6 and then cooled to 16°C – 20°C. Protein expression was induced with typically 250 μM IPTG and the cells were left over night at the latter temperature. The cells were collected by centrifugation at 4200 rpm for 25min at 4°C in a Beckman J6 centrifuge using a 6 x 1 L bucket rotor (4.2). The cells were resuspended in ice cold lysis buffer (50 mM Tris-HCl pH 7.5, 250 mM NaCl, 25 mM imidazole, 0.1 mM EGTA, 0.1 mM EDTA, 0.2 % Triton X-100, 10 μg/ml Leupeptin, 1 mM PefaBloc (Roche), 1mM DTT) and sonicated. Insoluble material was removed by centrifugation at 18500 xg for 25 min at 4°C. The supernatant was incubated for 1 h with Ni-NTA-agarose (Expedeon), then washed five times with 10 volumes of the lysis buffer and then twice in 50 mM HEPES pH 7.5, 150 mM NaCl, 0.015% Brij35, 1 mM DTT. Elution was achieved by incubation with the latter buffer containing 0.4M imidazole or by incubation with Tobacco Etch Virus (TEV) protease (purified in house). The proteins were buffer exchanged into 50 mM HEPES pH 7.5, 150 mM NaCl, 10% glycerol and 1 mM DTT and stored at -80°C. HOIP (697-1072) DU22629 and Itch (DU11097) ligases were expressed in BL21 cells as GST-tagged fusion proteins, purified via their tag and collected by elution (GST-Itch) or by removal of the GST-tag on the resin (HOIP).

### MDM2 E3 enzyme expression and purification

pGex-Mdm2 [DU 43570] was expressed in BL21 (DE3) *E. coli* cells grown in LB media containing 100 μg/ml ampicillin. Cells were induced with 250 μM isopropyl beta-D-1-thiogalactopyranoside (IPTG) at an OD600 of 0.6-0.8 and grown for 16 hours at 15°C. Cells were pelleted and resuspended in 50 mM Tris-HCl pH 7.5, 250 mM NaCl, 1 % Triton, 1 mM EDTA, 1 mM EGTA, 0.1 % 2-mercaptoethanol, 1 mM Pefabloc, 1 mM benzamidine. Cell lysis was carried out by sonication. After being clarified through centrifugation, bacterial lysate was incubated with Glutathione Agarose (Expedeon) for 2 hours at 4°C. The resin bound proteins were washed extensively with Wash buffer (50 mM Tris-HCl pH 7.5, 250 mM NaCl, 0.1 mM EGTA, 0.1 % 2-mercaptoethanol), before being eluted with wash buffer containing 20 mM Glutathione. The purified proteins were dialysed into storage buffer, flash frozen and stored at -80°C.

Plasmids generated at the University of Dundee for the present study are available to request on our reagents website (<u>https://mrcppureagents.dundee.ac.uk/</u>).

### E1/E2/E3 assay

The E2-E3 reaction consists of recombinant E1 (100 nM), E2 conjugating enzyme (125-250 nM), E3 ligases (250-500 nM) and 0.25 mg/mL BSA in 10 mM HEPES pH 8.5, 10 mM MgCl2 and 1 mM ATP in a total volume of 5 μl. Assays were performed by dispensing 2.5 μL of enzyme solution into round bottom 384-well plates (Greiner, Stonehouse, UK). Plates were centrifuged at 200 xg and the reactions were incubated at 37°C for 30 min. Reactions were initiated by the addition of 2.5 μL substrate solution containing 10 μM ubiquitin in 5 mM HEPES pH 8.5. For enzyme titration and time course experiments E2/E3 ligases concentrations ranged from 125 nM to 1000 nM with a maximum reaction time of 120 min. Plates were incubated at 37°C for typically 20-60 min (depending on the activity of E3 ligase used) before being quenched by the addition of 2.5 μL of a 10 % (v/v) trifluoroacetic acid (TFA) solution. Controls – with only DMSO - where placed on column 23. For the enzyme inactivated controls in columns 24, 2.5 μL of 10 % TFA was manually dispensed prior to addition of the enzyme solution by XRD-384 Automated Reagent Dispenser (FluidX). 1.05 μl of each reaction were spiked with 150 nl (4 μM) of ^15^N-labelled ubiquitin (average mass 8,659.3 Da) and 1.2 μl of 7.6 mg/ml 2,5-dihydroxyacetophenone (DHAP) matrix (prepared in 375 ml ethanol and 125 ml of an aqueous 25 mg/ml diammonium hydrogen citrate).

For E2 activity assays, we pre-incubated the Ube1 activating enzyme (100 nM) with 27 E2 conjugating enzymes at 1000 nM, 500 nM and 250 nM and stopped the reaction with 2% final TFA at different time points.

### Compound Library spotting and inhibitor screening

We used a library of 1430 FDA approved compounds from various commercial suppliers with validated biological and pharmacological activities. For single concentration screening 5 nL of 10 mM compound solution in DMSO was transferred into HiBase Low Binding 384-well flat bottom plates (Greiner bio-one) to give a final screening concentration of 10 μM. Columns 23 and 24 were reserved for uninhibited and inhibited controls respectively. The uninhibited control contained 5 nL DMSO but no compound, whereas the inhibited control contained 5 nL PR-619 but the enzyme was inactivated by pre-treatment with 1.0 % TFA. All compounds and DMSO were dispensed using an Echo acoustic dispenser (Labcyte, Sunnyvale, USA). For all HTS assays the final DMSO concentration was 0.1 %. For concentration response curves of known HOIP, MDM2 and ITCH inhibitors, a threefold serial dilution was prepared from 10 mM compound solutions in DMSO in 384-well base plates V-Bottom (Labtech). 100 nL of compound was transferred into 384-well round bottom low binding plates using a Mosquito Nanoliter pipetter (TTP Labtech, Melbourn, UK), giving a final concentration range between 100 μM and 100 nM.

### Target Spotting and MALDI mass spectrometry analysis

1,536-well AnchorChip MALDI targets (Bruker, Bremen Germany) were cleaned using 30% acetonitrile and 0.1% TFA and dried under a gentle flow of pure nitrogen. 200 nL matrix/assay mixture was spotted onto the AnchorChip Plates using a Mosquito nanoliter dispenser (TTP Labtech, Hertfordshire, UK). Spotted targets were air dried prior to MALDI TOF MS analysis. 0.25 μl of the resultant mixture was then spotted onto a 1,536 microtiter plate MALDI anchor target (Bruker, Bremen, Germany) using a Mosquito liquid handling robot (TTP Labtech, Melbourn, UK).

All samples were acquired on a Rapiflex MALDI TOF mass spectrometer (Bruker Daltonics, Bremen, Germany) high resolution MALDI TOF MS instrument with Compass for flexSeries 2.0. Reflector mode was used with optimized voltages for reflector 1 (20.82 kV), reflector 2 (1.085 kV) and reflector 3 (8.8 kV), ion sources (Ion Source 1: 20.0 kV, PIE 2.35 kV) and Pulsed Ion Extraction (500 ns). Matrix suppression has been set in Deflection mode to suppress matrix components up to 6560 *m/z*. Samples were run in automatic mode (AutoXecute, Bruker Daltonics) using the 1,536 spots AnchorChip. Ionization was achieved by a 10-kHz smartbeam-II solid state laser (run at 5 kHz) with Laser Fuzzy Control switched off, initial Laser Power set on “from Laser Attenuator” and accumulation parameters set to 4000 satisfactory shots in 500 shot steps. Movement parameters have been set on “Walk on Spot”. Spectra were accumulated by FlexControl software (version 4 Build 9), processed using FlexAnalysis software (version 4 Build 9) and the sophisticated numerical annotation procedure (‘SNAP’) peak detection algorithm, setting the signal-to-noise threshold at 5. Internal calibration was performed using the ubiquitin peak ([M+H]^+^ average = 8,659.3).

### Data analysis

A modified method for data acquisition was developed for FlexAnalysis Software version 4, using the SNAP algorithm. For area calculation, the complete isotopic distribution was taken into account. Data output was exported as a.csv file using FlexAnalysis series 4.0 (Build 24) Batch Process (Compass for flexseries 2.0). An in-house script has been used to extract - from the original csv output file - the data of interest. The script selects the area values of the light and the heavy ubiquitin and reports them in a grid with the same MALDI target geometry and sample positions within a txt file. Data are further analysed in Microsoft Excel, where plotting of graphs and IC50 calculation have been performed on Prism 7 software for windows, version 7.02.

For HTS data analysis, data was expressed as Activity % value for each test compound as follows:

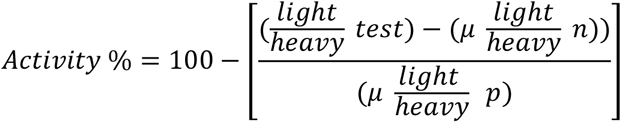

where 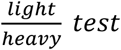 is the light ubiquitin signal normalized to the heavy ubiquitin signal associated with the test compound, 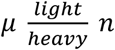 is the average of the light ubiquitin signal normalized to the heavy ubiquitin in the no inhibition signal positive controls (reaction in presence of DMSO only) and 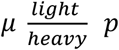 is the is the average of the light ubiquitin signal normalized to the heavy ubiquitin of maximum effect negative control wells (reaction where 2% TFA final has been added before the addition of substrate)

The performance of the assay on each screening plate was evaluated using internal controls to determine robust Z’ values, which were calculated as follows:

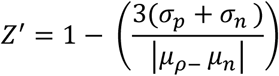

Where the means (*μ*) and standard deviations (*σ*) of both the positive (p) and negative (n) controls are reported.

### Author contributions

VDC performed all experiments; CJ, VB, AK and CJH produced proteins; VDC and MT designed experiments; VDC and MT prepared the manuscript with contributions from all authors.

## Acknowledgements

We would like to thank the DNA cloning, Protein Production, DNA sequencing facility and mass spectrometry teams of the MRC Protein Phosphorylation and Ubiquitylation Unit for their support. We would like to thank Prof Dario Alessi, Prof Ronald Hay, Prof Philip Cohen, Prof Katrin Rittinger, Dr. Sarah Buhrlage, Dr. Natalia Shpiro, Dr. Siddharth Bakshi, Dr. Andrea Testa and Dr. Francesca Morreale for tools and helpful discussions; Bruker Daltonics, particularly Meike Hamester, Rainer Paape and Anja Resemann for their technical support. We thank Dr. Anthony Hope, Alex Cookson and Lorna Campbell for providing the FDA approved compound library and support with the liquid handling. This work was funded by Medical Research Council UK (MC_UU_12016/5), the pharmaceutical companies supporting the Division of Signal Transduction Therapy (DSTT) (Boehringer-Ingelheim, GlaxoSmithKline and Merck KGaA).

## Supplementary Figures and Data

**Table S1:** Instantaneous reaction rate at 5 min of MDM2, ITCH and HOIP

| Instantaneous Rate at $t = 5$ min | | | | |
| --- | --- | --- | --- | --- |
| $\mu\text{M Ub}$ | 12.5 | 6.25 | 3.125 | |
| MDM2 | 0.61 | 0.24 | 0.30 | $\mu\text{mol/min}$ |
| ITCH | 0.41 | 0.34 | 0.30 | $\mu\text{mol/min}$ |
| HOIP | 1.85 | 1.28 | 0.94 | $\mu\text{mol/min}$ |

**Table S2:**
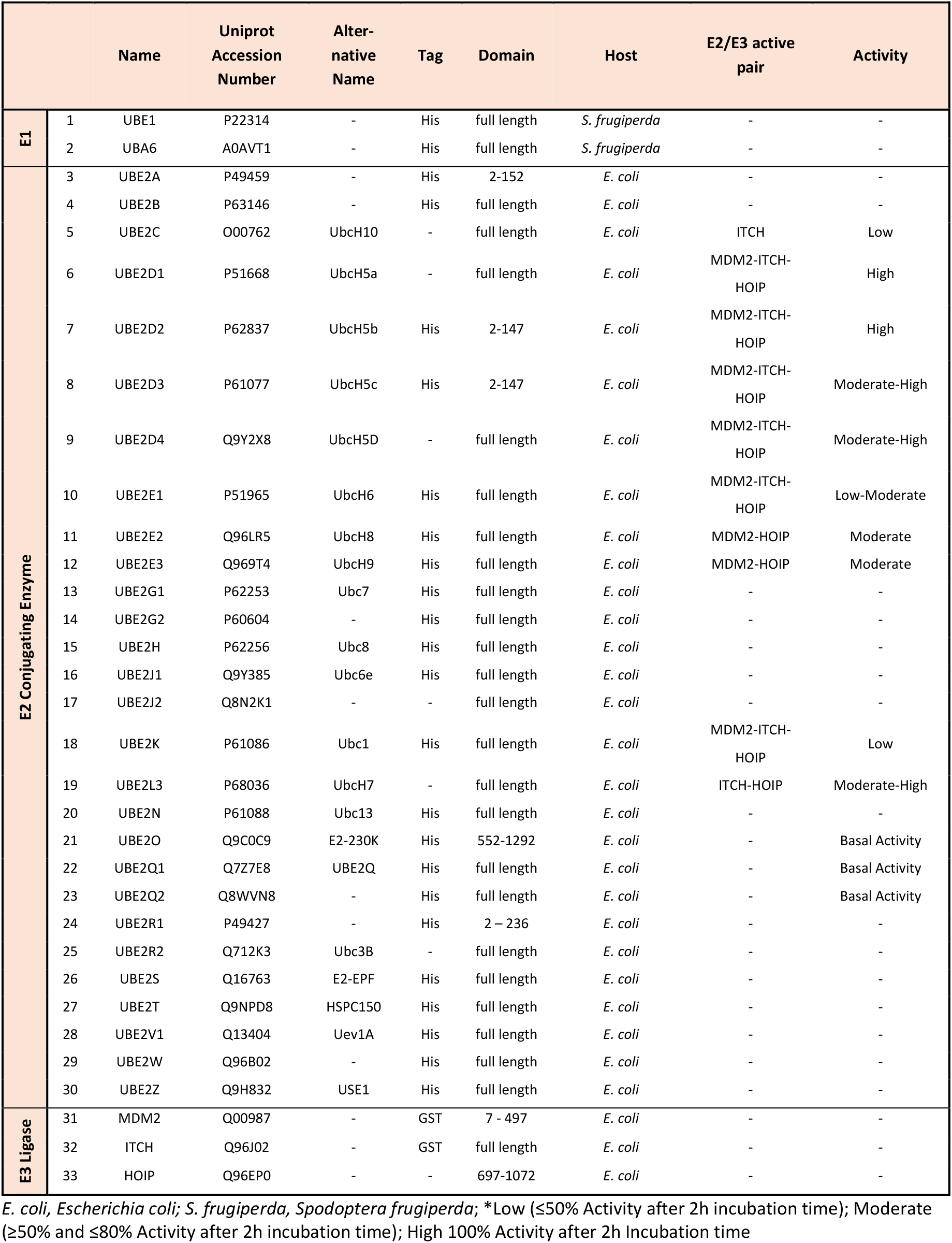
List of E1 Activating enzymes, E2 conjugating enzymes and E3 ligases used in this study

**Table S3|:** IC50 calculation of six E1, E2 or E3 inhibitors

| | IC50 ($\mu\text{M}$) | | | | | |
| --- | --- | --- | --- | --- | --- | --- |
|  | PR619 | Nutlin3A | Gliotoxin | Clomipramine | BAY117082 | PYR41 |
| MDM2 | 0.61 | Ambiguous | 0.50 | Ambiguous | Ambiguous | Ambiguous |
| ITCH | 0.41 | Ambiguous | 30.61 | 503.4 | 25.9 | 11.3 |
| HOIP | 0.22 | Ambiguous | 2.84 | Ambiguous | 2.9 | 5.7 |

**Table S4|:** Z-Prime scores

|  | Plate |  |  |  |  |
| --- | --- | --- | --- | --- | --- |
|  | A | B | C | D | E |
| MDM2 | 0.64 | 0.57 | 0.63 | 0.62 | 0.55 |
| ITCH | 0.67 | 0.64 | 0.73 | 0.83 | 0.79 |
| HOIP | 0.60 | 0.56 | 0.59 | 0.67 | 0.62 |

**Figure S1:**
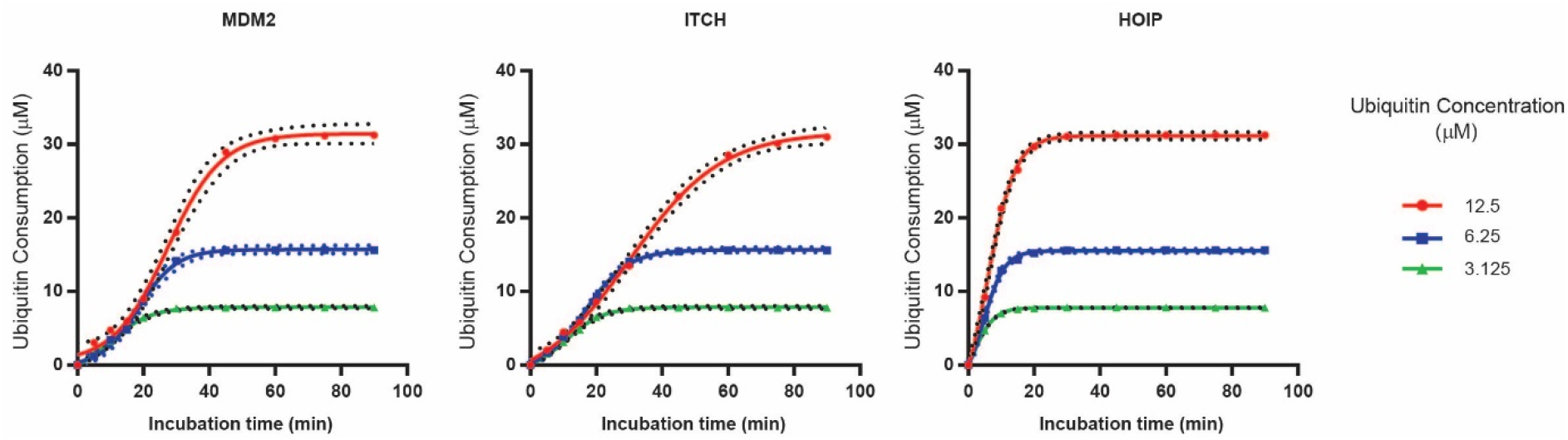
Substrate consumption over time of MDM2, ITCH and HOIP. Rate of disappearance of ubiquitin at different starting concentrations were measured for MDM2, ITCH and HOIP. Ubiquitin consumption (- (Ubiquitin μM t_2_ - Ubiquitin μM t_1_)) was plotted over time.

**Figure S2:**
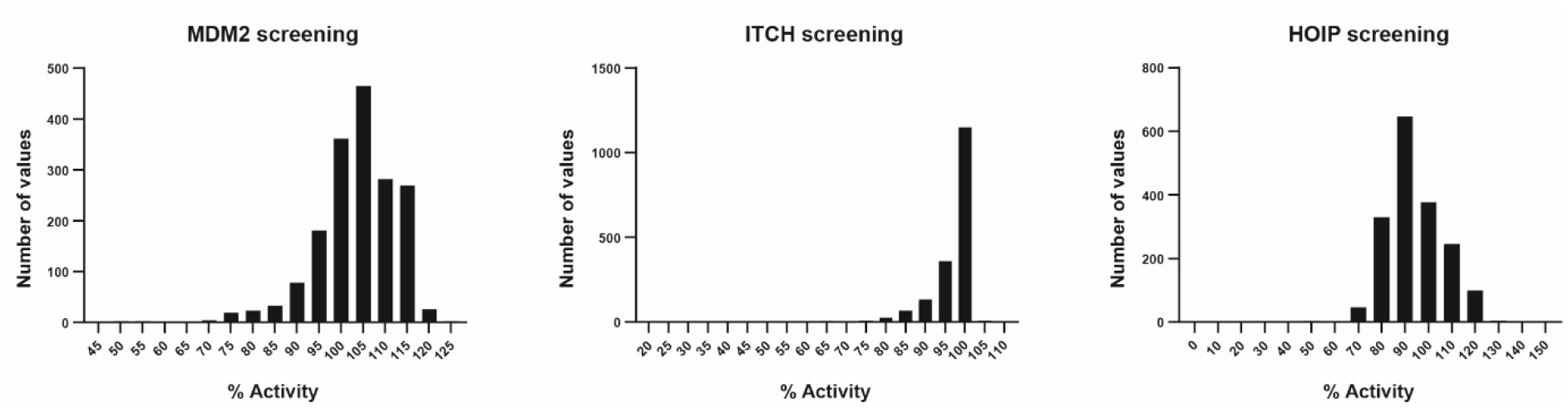
Frequency distribution of HTS data of MDM2, ITCH and HOIP. Bars report number of values falling within the reported % Activity.

**Figure S3:**
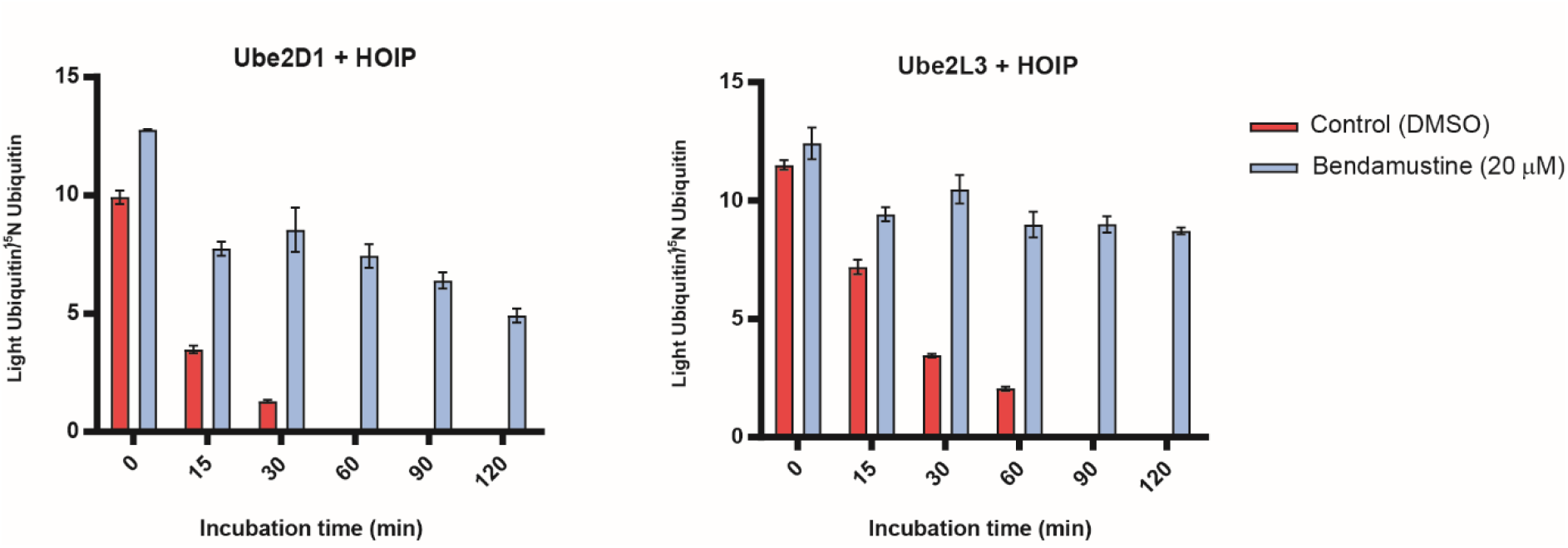
Bendamustine inhibits HOIP independently from the E2 conjugating enzyme. Bendamustine (20 μM) was tested against HOIP coupled with Ube2D1 and Ube2L3 over time. Bendamustine inhibits HOIP independently from the E2 in use.

